# TANGLED1 mediates interactions between microtubules that may promote spindle organization and phragmoplast guidance to the division site in maize

**DOI:** 10.1101/711796

**Authors:** Pablo Martinez, Ram Dixit, Rachappa S. Balkunde, Seán E. O’Leary, Kenneth A. Brakke, Carolyn G. Rasmussen

## Abstract

The microtubule cytoskeleton serves as a dynamic structural framework for mitosis in eukaryotic cells. TANGLED1 (TAN1) is a microtubule-binding protein that localizes to the division site and mitotic microtubules and plays a critical role in division plane orientation in plants. Here, in vitro experiments demonstrate that TAN1 directly binds microtubules, mediating microtubule zippering or end-on microtubule interactions, depending on their contact angle. Maize *tan1* mutant cells improperly position the preprophase band (PPB), which predicts the future division site. However, cell-shape-based modeling indicates that PPB positioning defects are likely a consequence of abnormal cell shapes and not due to TAN1 absence. Spindle defects in the *tan1* mutant suggest that TAN1-mediated microtubule zippering may contribute to metaphase spindle organization. In telophase, co-localization of growing microtubules ends from the phragmoplast with TAN1 at the division site suggests that TAN1 interacts with microtubule tips end-on. Together, our results suggest that TAN1 contributes to spindle and phragmoplast microtubule organization to ensure proper division plane orientation.

## Introduction

The proper organization of microtubule networks during interphase and mitosis is important to promote growth and development at both the cell and organismal levels (Wasteneys and Ambrose, 2009; Elliott and Shaw, 2018; Ehrhardt and Shaw, 2006; Baskin et al., 2004). Mechanisms for achieving and modulating microtubule organization are driven by microtubule-microtubule or microtubule-protein interactions: zippering at low contact angles (Ho et al., 2012; Tulin et al., 2012; Smertenko et al., 2004; Shaw et al., 2003), contact-mediated catastrophe (Dixit and Cyr, 2004), severing (Lindeboom et al., 2013; Zhang et al., 2013; Panteris et al., 2018; Komis et al., 2017) and stabilization at cell edges (Ambrose et al., 2011). These processes alter microtubule dynamics and organization. Mitotic microtubule structures are formed and modified by these activities to perform a distinct role in DNA segregation and separation of daughter cells. In plants, the key mitotic structures are the preprophase band (PPB), metaphase spindle, and phragmoplast. Proteins which regulate the formation and function of these structures are localized along these different structures as well as the cortical plant division site.

During the G2 phase of the cell cycle, the preprophase band (PPB) is formed as a ring-shaped arrangement of microtubules, actin and associated proteins that localize just under the plasma membrane to form the cortical division zone (Smertenko et al., 2017; Van Damme et al., 2007). The PPB is an early marker of the future division site in land plants: it indicates the location where the developing new cell wall will fuse with the mother cell (Rasmussen and Bellinger, 2018; Facette et al., 2019; Pickett-Heaps and Northcote, 1966). Several microtubule associated proteins play an important role in division plane orientation by promoting PPB formation. A large family of proteins with microtubule binding motifs recruit a protein phosphatase type 2A (PP2A) complex to form the PPB (Spinner et al., 2013; Wright et al., 2009; Traas et al., 1995; Spinner et al., 2010; Drevensek et al., 2012; Schaefer et al., 2017). The proper formation and positioning of the PPB may orient the metaphase spindle to promote rapid mitotic progression (Chan et al., 2005; Ambrose and Cyr, 2008; Schaefer et al., 2017). As cells enter metaphase, the PPB is completely disassembled; however a handful of proteins that colocalize with the PPB continue to label the division site until the end of cytokinesis (Walker et al., 2007; Xu et al., 2008; Lipka et al., 2014; Martinez et al., 2017; Li et al., 2017; Buschmann et al., 2015).

During telophase, the phragmoplast is assembled from microtubules, actin, and associated proteins to aid in the formation of the cell plate via vesicle delivery (Smertenko et al., 2017; Smertenko, 2018; Lee and Liu, 2013; Jürgens, 2005b). The phragmoplast expands outwards to the cell cortex through the polymerization of new microtubules from existing leading edge microtubules and depolymerization at the lagging edge as the cell plate is assembled (Murata et al., 2013). The direction of phragmoplast expansion is thought to be guided by proteins that continuously label the division site (Rasmussen and Bellinger, 2018; Livanos and Müller, 2019). Once the phragmoplast reaches the cortex it is disassembled and the cell plate fuses with the plasma membrane, completing cytokinesis (Jürgens, 2005a; Worden et al., 2012). Mutants with defects in maintaining division plane orientation place new cell walls outside the location originally specified by the PPB. In maize, *tangled1* (*tan1)* mutants have division plane defects in both symmetric and asymmetric divisions (Smith et al., 1996) caused by a failure of the phragmoplast to return to the division site originally indicated by the PPB (Martinez et al., 2017). TAN1-YFP localizes to the cortical division site throughout mitosis in *Arabidopsis* and maize (Martinez et al., 2017; Walker et al., 2007). TAN1 also co-localizes with mitotic microtubule arrays in vivo when fused to YFP (Martinez et al., 2017) and using a non-specific TAN1 antibody (Smith et al., 2001).

Double mutants for two kinesin 12 paralogs in *Arabidopsis thaliana, phragmoplast orienting kinesin 1 (pok1)* and *pok2* display a severe division plane defect (Müller et al., 2006b). POK1 interacts directly with TAN1 and localizes to the division site (Walker et al., 2007; Lipka et al., 2014; Rasmussen et al., 2011). Both POK1 and POK2 are required for TAN1 localization to the division site after metaphase (Walker et al., 2007; Lipka et al., 2014). POK2 acts as a weak microtubule plus-end-directed motor in vitro (Chugh et al., 2018). Interestingly, in addition to its division site localization, POK2 also accumulates in the phragmoplast midline where it may interact with MICROTUBULE ASSOCIATED PROTEIN65-3, MAP65-3, or other MAP65 proteins (Herrmann et al., 2018; Ho et al., 2011). Another closely related MAP65, MAP65-4, is localized to the PPB, spindle and phragmoplast and the division site (Li et al., 2017). The *map65-3 map65-4* double mutant in *Arabidopsis thaliana* displays a severe cytokinesis defect but it is not yet clear whether it has a division plane defect (Li et al., 2017). MAP65-4 regulates microtubule stability by increasing microtubule elongation phases during bundling (Fache et al., 2010) while another related MAP65, MAP65-1 increases microtubule stability by protecting against severing and promoting microtubule flexibility during bundling (Portran et al., 2013; Stoppin-Mellet et al., 2013; Burkart and Dixit, 2019). Microtubule binding and bundling proteins therefore may contribute to the assembly of the mitotic microtubule structures, but also serve as important effectors for the establishment, timely progression and execution of properly oriented plant cell divisions.

In addition to division plane defects, the *tan1* mutant has mitotic progression delays and reduced plant stature (Martinez et al., 2017; Smith et al., 1996). Mitotic progression delays and phragmoplast guidance defects were mostly uncoupled using a partially rescued *tan1* mutant expressing TAN1-YFP fused to the CYCLIN B-destruction box motif (Martinez et al., 2017). In this partially rescued line, mitotic delays are observed but division plane defects are rare, coinciding with TAN1-YFP signal at the division site, but lack of detectable TAN1-YFP signal in the spindle and phragmoplast. We hypothesize that TAN1 is a multifunctional protein that aids in timely mitotic progression when it localizes to mitotic microtubule structures and maintains division plane orientation via phragmoplast guidance when it is localized to the division site. Here we report an in vitro function for TAN1 in mediating microtubule interactions, and an in vivo function in spindle organization and phragmoplast microtubule interactions at the division site.

## Results and Discussion

### TAN1 binds to microtubules in vitro

TAN1 protein has been shown to bind to taxol-stabilized microtubules in a blot overlay assay (Smith et al., 2001). To quantitatively assess the binding of TAN1 to microtubules, we recombinantly expressed 6xHIS-tagged ZmTAN1 (HIS-TAN1) protein. HIS-TAN1 protein bound to taxol-stabilized microtubules in cosedimentation experiments (Figure 1A). By titrating the microtubule concentration against a fixed concentration of HIS-TAN1, we calculated an apparent equilibrium dissociation constant of 1.27μM (Figure 1B). This affinity is similar to that of other microtubule-binding proteins (Tulin et al., 2012; Portran et al., 2013; Wong and Hashimoto, 2017). The precise molecular mechanism of TAN1-microtubule interaction is unknown, therefore the apparent affinity may be a composite of distinct microtubule binding modes. To directly visualize the binding of TAN1 to microtubules in vitro, we purified recombinant HIS-TAN1-GFP. Unfortunately, this fusion protein was not fluorescent, probably because GFP did not fold correctly during renaturation of recombinant protein from bacterial inclusion bodies. Since HIS-TAN1-GFP still bound to microtubules (Supplemental Figure 1A), we labeled it with the organic fluorophore Atto488 to visualize it using fluorescence microscopy. When co-incubated with taxol-stabilized rhodamine-labeled microtubules, Atto488-tagged HIS-TAN1-GFP localized relatively uniformly along the microtubule lattice (Figure 1D-E).

**Figure 1:**
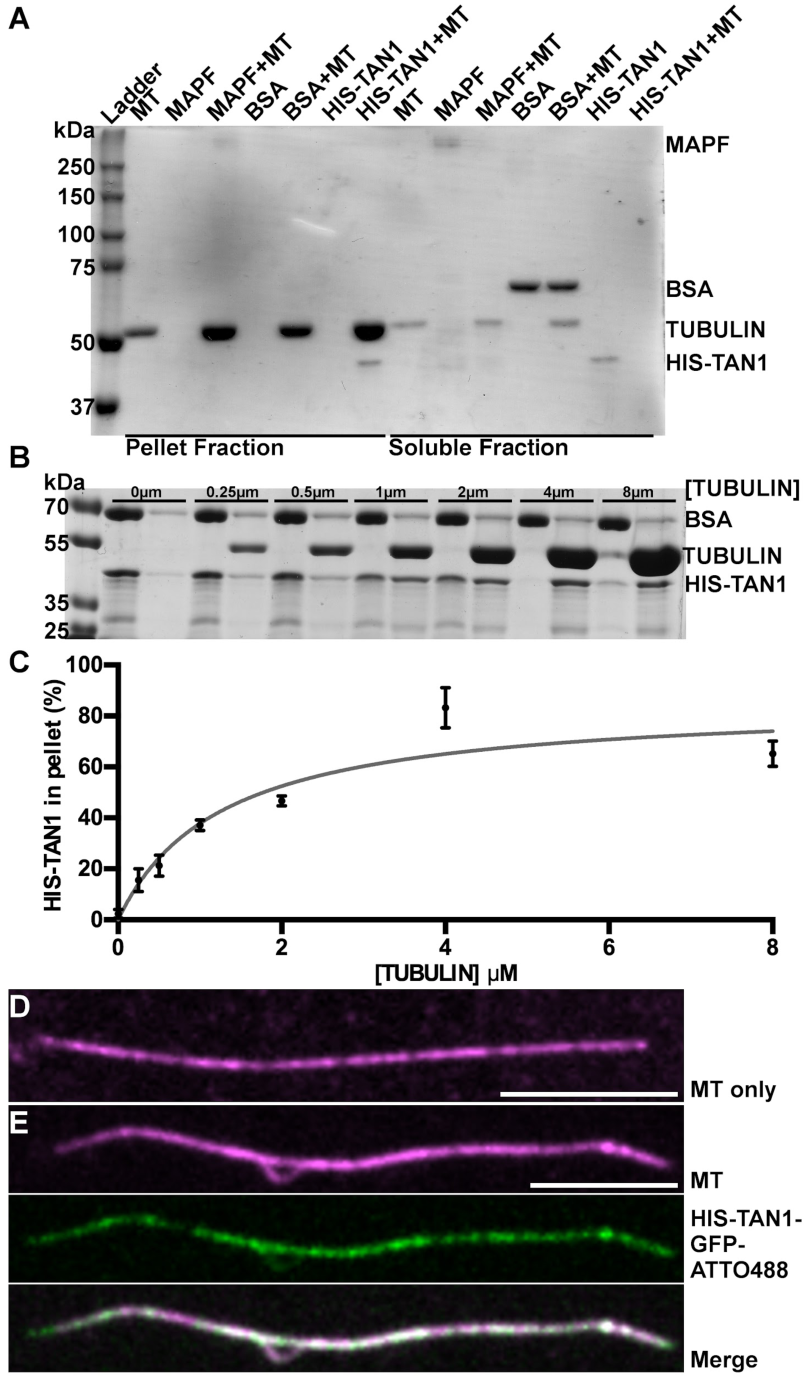
Recombinantly expressed TAN1 binds to microtubules. (A) Coomassie stained SDS PAGE results from microtubule cosedimentation with HIS-TAN1, positive control (MAPF, 70% MAP2) and negative control (BSA) controls separated into corresponding pellet and soluble fractions. (B) Coomassie stained SDS PAGE results from microtubule cosedimentation assay using 2μM HIS-TAN1 and from 0-8μM of tubulin. (C) Hyperbolic fit of microtubule cosedimentation data for HIS-TAN1 at varying concentrations of microtubules determines an apparent equilibrium dissociation constant of 1.27μM +/- 0.3 (S.D.). (D, E) Rhodamine-labeled, taxol-stabilized microtubule in buffer only control (magenta) (D) or incubated with HIS-TAN1-GFP-Atto488 (green). Scale bar is 10μm.

### TAN1 does not markedly alter microtubule dynamics in vitro

To determine whether TAN1 regulates microtubule polymerization dynamics, we conducted in vitro microtubule polymerization experiments on coverslips. Microtubules were nucleated from GMPCPP-stabilized microtubule seeds and their polymerization and depolymerization was visualized using rhodamine-labeled tubulin and total internal reflection fluorescence microscopy (materials and methods). At relatively low concentrations of HIS-TAN1 (< 1μM), no significant effect on microtubule dynamics was observed (Table 1). At a concentration of 2 μM HIS-TAN1, which is close to the apparent K_d_ of TAN1 for microtubules, we observed small decreases in both microtubule plus-end growth and plus-end shrinkage rates (Table 1). HIS-TAN1 addition did not alter the amount of time microtubules spent growing or the frequency of catastrophes. However, small but significant differences in time spent shrinking were observed (Table 1). Under the experimental conditions used, rescue events were rare and the minus-ends were not dynamic; therefore, these parameters were not quantified. Together, these results suggest that regulation of microtubule polymerization dynamics is unlikely to be the primary function of TAN1.

**Table 1:** Summary of microtubule dynamics and microtubule interactions at different concentrations of HIS-TAN1. Significance indicated by (*) p-value > 0.05, (**) p-value > 0.01, (***) p-value > 0.001.

| Plus-end dynamics | 0μM HIS-TAN1 | 0.1 μM HIS-TAN1 | 0.5 μM HIS-TAN1 | 1μM HIS-TAN1 | 2μM HIS-TAN1 |
| --- | --- | --- | --- | --- | --- |
| Growth events (n) | 156 | 180 | 166 | 214 | 196 |
| Growth Rate (μm/sec, mean +/- S.D.) | 1.8 +/- 0.4 | 1.8 +/- 0.3 | 1.8 +/- 0.3 | *1.6 +/- 0.5 | ***1.5 +/- 0.3 |
| Shrinkage events (n) | 109 | 127 | 113 | 153 | 149 |
| Shrinkage Rate (μm/sec, mean +/- S.D.) | 31.5 +/- 15.6 | 27.7 +/- 10.8 | *26.2 +/- 8.8 | 27.8 +/- 9.7 | ***22 +/- 10.0 |
| Time growing (%) | 94.9 | 93.8 | 94.5 | 94.6 | 95.2 |
| Time shrinking (%) | 5.1 | **6.2 | *5.5 | 5.4 | 4.8 |
| Catastrophe Frequency (events/minute) | 0.3 | 0.4 | 0.4 | 0.4 | 0.5 |
| Crossovers (n) | 445 | 346 | 334 | 334 | 506 |
| Bundling events (n) | 2 | 0 | 0 | 3 | 139 |
| Bundling frequency (%) | 0.5 | 0 | 0 | 0.9 | 27.5 |

### HIS-TAN1 mediates lateral and end-on microtubule interactions in vitro

During the course of our in vitro microtubule dynamics experiments, we observed that 2μM HIS-TAN1 mediated interactions between microtubules that happened to contact each other. To promote microtubule interactions, we conducted these next experiments with a higher concentration of GMPCPP-stabilized seeds and free tubulin dimers to produce more microtubules that grew longer and hence encountered each other more frequently. We used 2μM HIS-TAN1 because it results in microtubule interactions (139 interaction events resulting from 506 crossovers), whereas none are observed at lower concentrations of HIS-TAN1 (Table 1). We observed two kinds of microtubule interactions depending on the microtubule contact angle. At small or shallow contact angles (angle = 19.6° +/- 7.6° average +/- S.D.), the microtubules progressively zippered together to produce bundles (n = 47 bundling events out of a total of 139 interactions observed, 34% of bundling events) (Figure 2A-2B). Zippering of microtubules in parallel and antiparallel configurations occurred with similar frequencies (n = 13/27 and 14/27 where orientation was unambiguous, respectively). Therefore, TAN1 does not preferentially bundle microtubules in specific orientations in contrast to the MAP65 microtubule bundling proteins, that preferentially bundle antiparallel microtubules. (Gaillard et al., 2008; Tulin et al., 2012). At high contact angles (angle = 60° +/- 20° average +/- S.D.), transient “end-on” microtubule interactions were observed during microtubule depolymerization (Figure 2C-2D, Supplemental Video 1). As one microtubule depolymerized past a previous crossover site, TAN1 mediated an interaction at the crossover point, the depolymerizing end stayed bound to the sidewall of the second microtubule, resulting in a pulling force on the stable microtubule (n = 92 end-on interactions out of a total of 139 interactions observed, 66% of interaction events). Based on these data, we conclude that the outcome of TAN1-microtubule interactions depend on the initial contact or crossover angle between the microtubules, and that at high contact angles, TAN1-microtubule interactions lead to transient pulling or catching.

**Figure 2:**
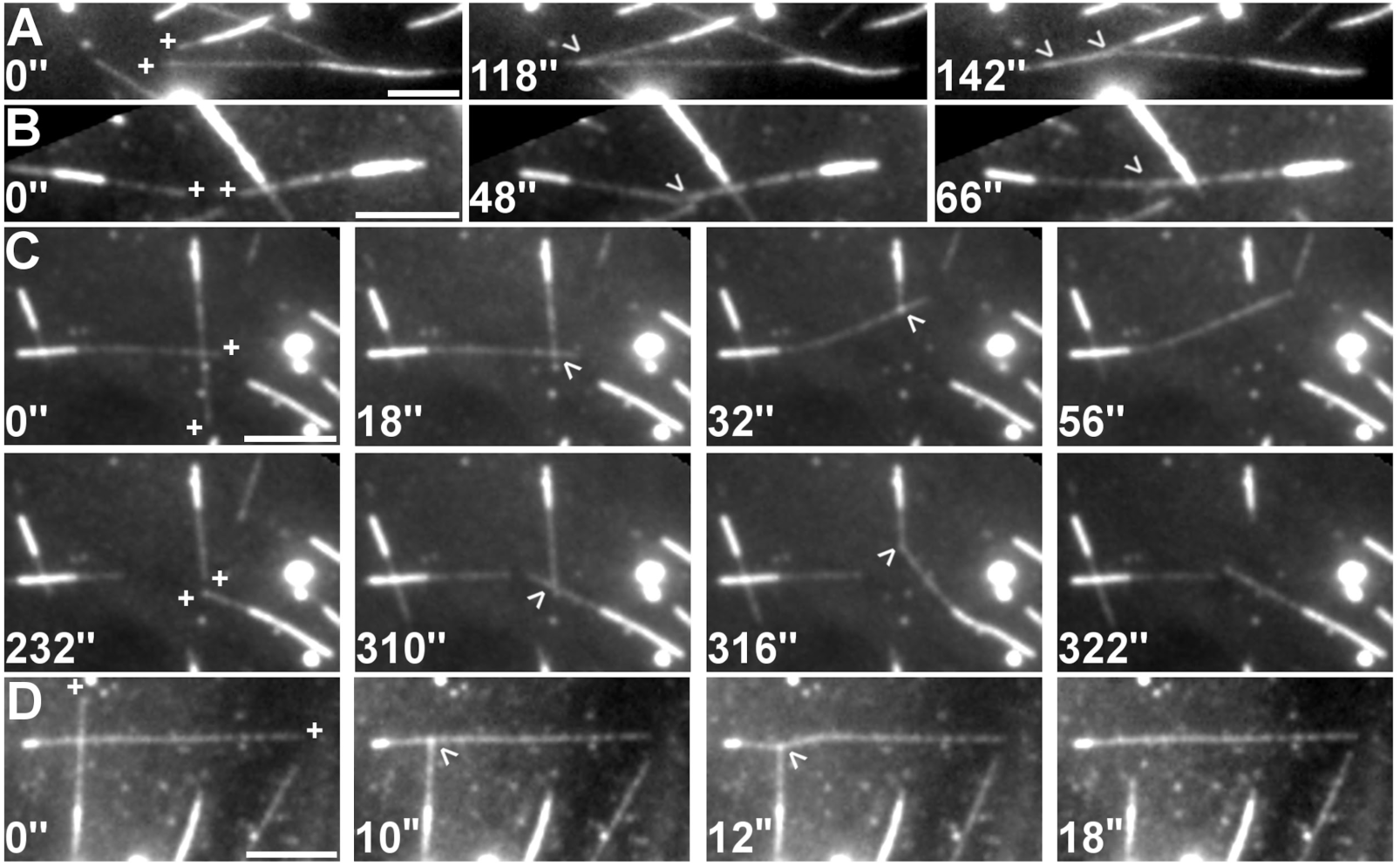
HIS-TAN1 contact-angle-independent dynamic microtubule interactions. (A-D) Dynamic rhodamine-labelled microtubules nucleated from GMPCPP-stabilized seeds with plus-ends indicated by a (+) and a crossover indicated with an arrowhead. Microtubule seeds are identified by their brighter signal compared to the growing microtubule ends. 2μM HIS-TAN1 is present in the assay. (A) Two microtubule plus-ends are indicated with their plus-ends polymerizing in the same direction. These microtubules encounter each other in a parallel orientation and are zippered together. (B) Two microtubule plus-ends are indicated with their plus-ends growing towards each other. These microtubules are zippered together in an antiparallel orientation. (C) Two microtubule plus-ends are indicated at the start (0”). These microtubules crossover and at 18” one of them depolymerizes. The depolymerizing end of this microtubule appears to pull on the other microtubule over the course of depolymerization. At 232” (new plus-end growth indicated) a new crossover is formed followed by a depolymerization event which again pulls at the crossover with the non-depolymerizing microtubule (316”). Timelapse shown in Supplemental Video 1. (D) Two microtubule plus-ends are indicated at the start (0”) which cross over at a high angle (∼90°). Depolymerization of one microtubule leads to transient deformation of the other microtubule at the crossover point. Scale bar is 10μm.

Microtubule zippering is a well-characterized form of microtubule bundling in plants, animals and fungi (Dixit and Cyr, 2004; Tulin et al., 2012; Janson et al., 2007; Subramanian et al., 2010; Gaillard et al., 2008). Microtubule end-on interactions have been studied extensively in animals and fungi and typically involve forces generated by motor proteins (Laan et al., 2012b; a). For example, end-on microtubule capture by motor proteins is important for spindle positioning in animals (Kiyomitsu, 2019) and yeast (Gupta et al., 2006). Non-motor dependent mechanisms, such as harnessing the energy of a depolymerizing microtubule, also generate pulling forces (Dogterom et al., 2005; Grishchuk et al., 2005). TAN1, because it lacks canonical motor domains, is unlikely to be a motor protein. Currently, we do not know how many TAN1 molecules are required for effective microtubule end-on interactions or whether the strength of these interactions is a function of the number of TAN1 molecules.

### Abnormal cell shape is likely responsible for spatial positioning defects of the PPB in the *tan1* mutant

Previous work indicated that the orientation of the PPB is more variable in *tan1* mutants compared to WT, indicative of a PPB placement defect (Cleary and Smith, 1998; Mir et al., 2018). However, these measurements were obtained from 2D micrographs of PPBs and might not accurately reflect the proper placement of the PPB in 3D, particularly in cells with irregular shapes. To overcome this shortcoming, we collected confocal Z-stacks and used MorphoGraphX (Barbier de Reuille et al., 2015) to extract wild-type and *tan1* mutant three-dimensional cell shapes (Figure 3A-B). We then used Surface Evolver to predict division planes in these cells using a mathematical modeling approach (Martinez et al., 2018; Brakke, 1992). The predicted division planes were then compared to the in vivo PPB location (Figure 3A-B). For wild-type cells, the average PPB offset from the predicted divisions was 0.40μm^2^ +/- 0.96 (average +/- standard deviation (SD) n = 16), while PPB offset was higher in *tan1* mutants (PPB offset = 1.85μm^2^ +/- 3.93, average +/- S.D. n=45; p-value = 0.0012 Mann-Whitney, Figure 3C). These data suggest that TAN1 might be required for proper PPB placement. Alternatively, the PPB placement defects may be an indirect consequence of abnormal cell shapes in the *tan1* mutant. Therefore, we developed a quantitative method to compare cell shapes called the “abnormality index” by measuring the deviation from the surface area center and volume center (see Materials and Methods). WT cells had lower and more consistent abnormality index compared to *tan1* mutant cells (Figure 3D, WT n=16 abnormality index = 0.14 +/- 0.1, *tan1* n=45 abnormality index = 0.37 +/- 0.32 p-value = < 0.0001 Mann-Whitney; Average +/- S.D.). These data confirm that wild-type plants tend to have normally shaped cells, while *tan1* mutants have cells with both normal and abnormal shapes, consistent with our observations.

**Figure 3:**
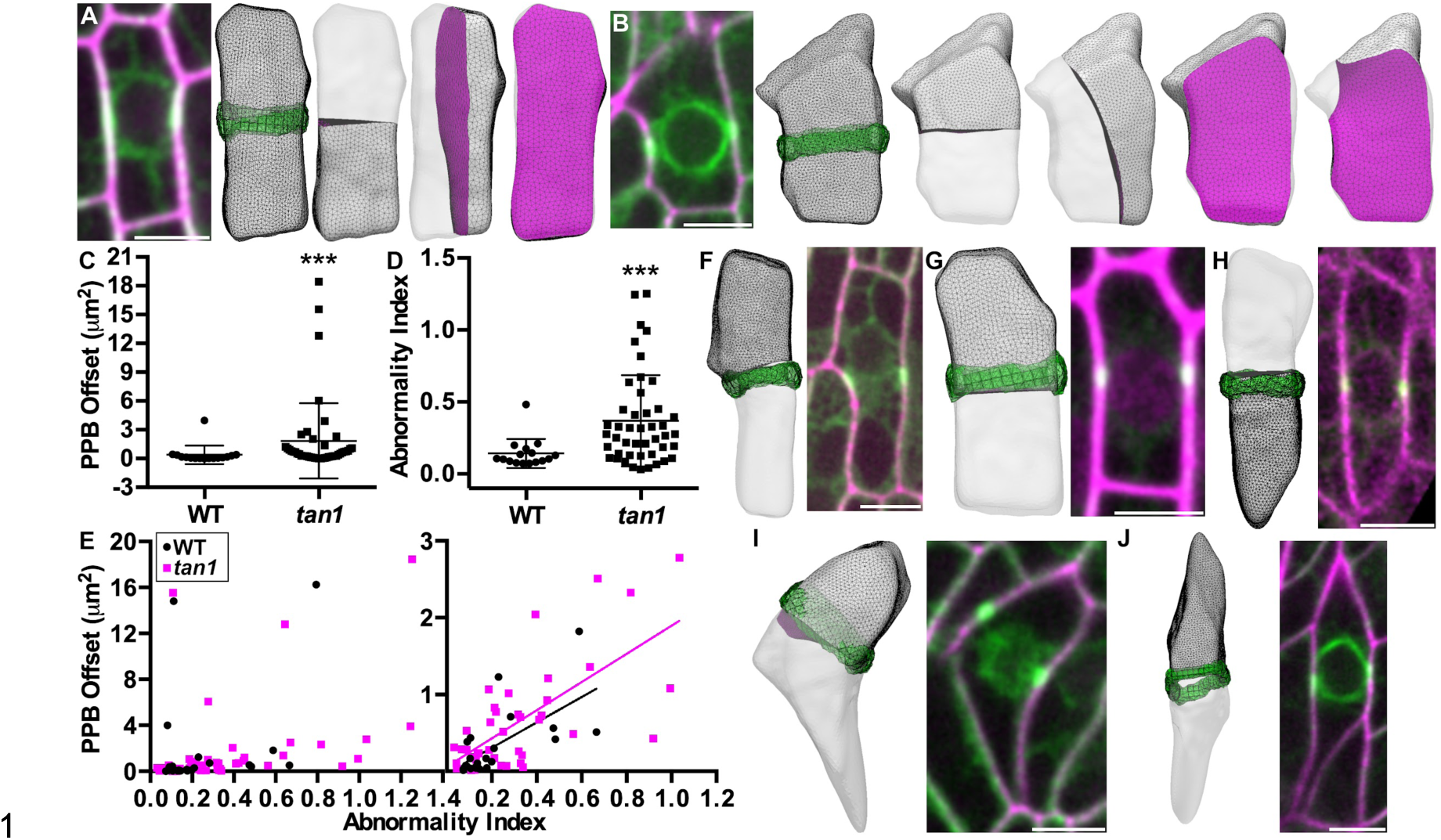
Abnormally shaped cells have higher PPB offset in wild-type and *tan1* cells. (A) Micrograph of wild-type maize leaf epidermal cell expressing YFP-TUBULIN (green) stained with propidium iodide (magenta) next to the 3D cell shape reconstruction smoothed with 30th degree spherical harmonics overlaid with PPB (green) and predicted (left to right) transverse, longitudinal, periclinal division planes (magenta) overlaid. (B) Micrograph of *tan1* maize leaf epidermal cell next to cell shape reconstruction with PPB overlaid and (from left to right) transverse, longitudinal, periclinal and other division planes. (C) PPB offset in *tan1* mutants is significantly higher than WT (WT n=16 PPB offset = 0.40μm^2^ +/- 0.96, *tan1* n=45 1.85μm^2^ +/- 3.93, average +/- S.D.; p-value = 0.0012 (Mann-Whitney). (D) Abnormality index in *tan1* mutants is significantly higher compared to WT (WT n=16 abnormality index = 0.14 +/- 0.1, *tan1* n=45 abnormality index = 0.37 +/- 0.32 p-value = < 0.0001 Mann-Whitney; Average +/- S.D.) (E) Abnormality index versus PPB offset suggests abnormal cell shapes generally show higher offsets, WT r = 0.53, p value = .007 and *tan1* r = 0.58, p value <0.0001 (Spearman correlation coefficient). A subset of data (values of PPB offset < 3) is displayed adjacent with a linear fit of WT r = 0.7, p value = 0.0003, n = 22 and *tan1* r = 0.6, p value = < 0.0001, n = 40 (Spearman correlation coefficient). (F-J) Best-fit predicted divisions overlaid with in vivo PPB location next to corresponding micrograph of maize epidermal cells expressing YFP-TUBULIN (green) and either expressing membrane marker PIP2-CFP (H, magenta) or stained with propidium iodide (F-G, I-J, magenta) to outline the cell shape. (F) Example of a wild-type cell with abnormality index of 0.59 and PPB offset of 1.82μm^2^. (G) Example of a wild-type cell with abnormality index of 0.38 and PPB offset of 0.09μm^2^. (H) Example of a *tan1* mutant cell with abnormality index of 0.32 and PPB offset of 0.26μm^2^. (I) Example of a *tan1* mutant cell with abnormality index of 1.25 and PPB offset of 3.92μm^2^. (J) Example *tan1* mutant cell with abnormality index of 0.99 and PPB offset of 1.08μm^2^. Scale bar is 10μm.

If TAN1 plays a direct role in PPB placement, we would expect abnormal PPB placement in *tan1* mutants regardless of cell shape. In contrast, we found a positive correlation between abnormality index and PPB offset in *tan1* mutant cells (Spearman correlation coefficient = 0.58, p value = <0.0001, n=45 cells), suggesting that PPB placement deviated from predicted divisions more strongly in highly abnormal cells shapes. To address whether this trend was similar in wild-type cells, we specifically looked for and modeled additional wild-type cells which displayed altered cell shapes with high abnormality indices (Spearman correlation coefficient = 0.53, p value = 0.007 n = 25 cells). Both wild-type and *tan1* mutant cells with higher abnormality indices typically had higher PPB offsets for the whole dataset (Figure 3E, left panel) as well as the dataset removing outliers (Figure 3E, right panel), with examples of cells shown in (Figure 3F-J). Due to the correlation between PPB placement defects and aberrant cell shapes in *tan1* mutants, we hypothesize that defects in PPB placement are a consequence of cell shape abnormalities and not directly related to TAN1 function during G2. Modeling approaches based on microtubule organization suggest that interphase cortical microtubule arrangements may be an important modulator in PPB positioning (Chakrabortty et al., 2018; Mirabet et al., 2018). The orientation of the PPB typically follows the orientation of the prior interphase microtubule array (Flanders et al., 1989; Gunning and Sammut, 1990) Our result suggests that intrinsically abnormally shaped cells may lead, in the next round of cell division, toward less geometrically accurately placed PPBs. This effect may explain why other division plane mutants have offset or oblique PPBs (Pietra et al., 2013; Müller et al., 2006a). Additionally, mutants with cell expansion defects that cause aberrant cell shapes may also lead first to misoriented PPBs and then apparent division plane defects.

### Spindle organization is disrupted in the *tan1* mutant

Previously, we showed that *tan1* mutant cells had mitotic progression delays during metaphase and telophase, but no specific hypothesis was proposed to explain why delays occurred (Martinez et al., 2017). If TAN1 plays a significant role in crosslinking spindle microtubules, metaphase delays may reflect defective spindle organization. Using time-lapse imaging, we assessed overall spindle morphology in maize leaf cells expressing YFP-TUBULIN. In WT cells, we always observed bipolar spindles (n = 38) (Figure 4A). In *tan1* mutant cells, spindles were occasionally collapsed or misorganized (13.5% n = 5/35), but later recovered into typical bipolar spindles (Figure 4B). Metaphase delays previously described in *tan1* mutants occurred frequently, leading to an average 1.5x time delay compared to wild-type (Martinez et al., 2017), whereas spindle morphology defects were more rare. This suggests that defects in microtubule organization only occasionally lead to detectable spindle collapse in the *tan1* mutant, consistent with redundant mechanisms for spindle assembly. Bundling of metaphase spindle microtubules is important for proper and timely spindle assembly (Masoud et al., 2013; Mullen and Wignall, 2017; Ambrose and Cyr, 2007; Winters et al., 2019). Based on in vitro microtubule zippering by TAN1, it is likely that TAN1 mediates bundling of spindle microtubules as they encounter each other at shallow angles. Thus, TAN1 localization to the spindle might be important for correct spindle assembly and mitotic progression through metaphase.

**Figure 4:**
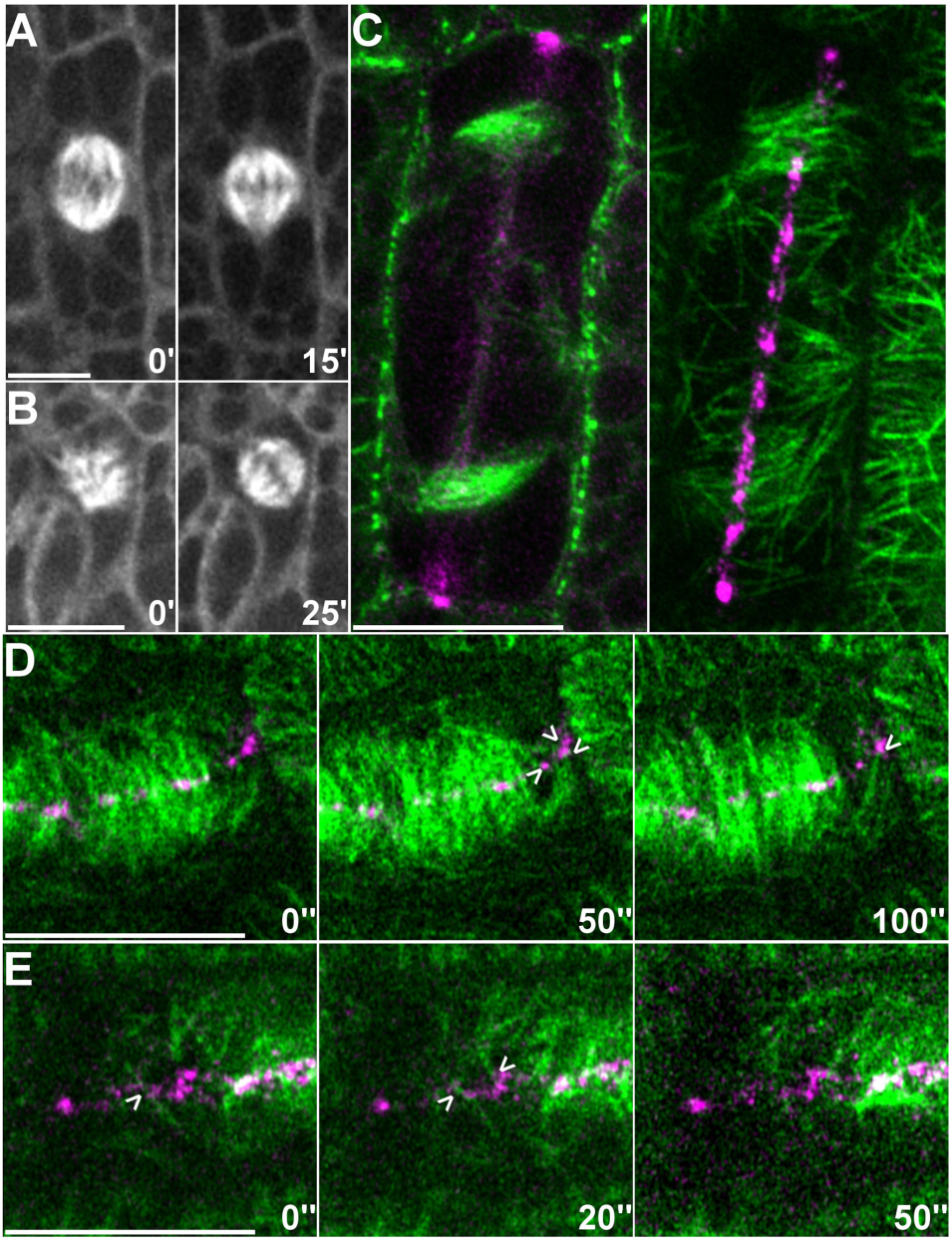
TAN1-mediated microtubule interactions may organize spindles and promote proper phragmoplast guidance. (A) Wild-type maize epidermal cell expressing YFP-TUBULIN displays normal bipolar spindle morphology over time. (B) *tan1* maize epidermal cell expressing YFP-TUBULIN which displays a misorganized spindle that recovers to a canonical bipolar organization. (C-D) Wild-type maize epidermal cells expressing CFP-TUBULIN (green) and TAN1-YFP (magenta). (C) Micrographs display both at the midplane and cortex of cell undergoing a longitudinal division. Phragmoplast and leading edge microtubules at the cortex are positioned at the division site and are partially colocalized with TAN1. (D-E) Timelapse showing potential phragmoplast leading edge microtubule contact and interaction with TAN1 at the division site (arrowheads). Figure 4D is shown in Supplemental Video 2. Scale bar is 10μm.

### Microtubules and TAN1 co-localize at the division site during telophase

To understand how TAN1 might mediate phragmoplast guidance during telophase (Martinez et al., 2017; Mir et al., 2018), we imaged TAN1 and microtubules at the division site. CFP-TUBULIN labeled microtubules and TAN1-YFP were imaged in cells undergoing longitudinal divisions, where phragmoplast guidance is more readily visualized. Colocalization of CFP-TUBULIN and TAN1-YFP at the division site was assessed at the cell cortex after initial phragmoplast contact. A small number of phragmoplast microtubules co-localize with TAN1 puncta (Pearson’s correlation coefficient 0.23 +/- 0.078 avg +/- S.D., n = 21), but about half of the TAN1 puncta were associated with microtubules (Manders overlay coefficient, C = 0.41 +/- 0.1 avg +/- S.D, Figure 4C). Together, these results suggest that a subpopulation of microtubules from the leading edge of the phragmoplast interact with cortical TAN1 puncta as the phragmoplast expands across the division site (Figure 4D-E, Supplemental Video 2).

Models for phragmoplast guidance previously proposed that leading edge phragmoplast microtubules interact with proteins at the cortical division site either through specific protein-protein interactions or microtubule-protein interactions (Herrmann et al., 2018; Lipka et al., 2014; Li et al., 2017). POK2, which is localized to the division site, was shown to be a plus-end directed kinesin (Chugh et al., 2018). POK2 may effectively push against the plus-ends of microtubules which encounter the division site (Chugh et al., 2018). POK2 also directly interacts with MAP65-3, which is localized to bundled microtubules both at the phragmoplast midzone and leading edge, serving as another potential type of interaction between the phragmoplast and the division site. The localization of TAN1 at the division site is important for its function in phragmoplast guidance (Mir et al., 2018; Martinez et al., 2017). Based on the results from this study, we propose that end-on interactions between the plus-ends of phragmoplast leading edge microtubules and TAN1-YFP puncta at the division site serve to exert pulling forces on these microtubules to guide phragmoplast trajectory.

While TAN1 has long been characterized as a microtubule binding protein, the functional significance of this finding remained elusive. Our in vitro analysis of TAN1-microtubule activities combined with live-imaging observations of TAN1 localization on spindle microtubules and at phragmoplast leading edge microtubule tips suggest that TAN1 leads to different outcomes depending on the geometry of microtubule encounters. This provides a plausible explanation for how TAN1 contributes to both spindle formation and phragmoplast guidance.

## Materials and Methods

### HIS-TAN1 and HIS-TAN1-GFP purification and labeling

A codon-optimized cDNA encoding the maize HIS-TAN1 and HIS-TAN1-GFP was synthesized in vitro, followed by protein expression and purification, all performed by Genscript (Genscript Corp Piscataway, New Jersey USA). *E. coli* strain SHuffle was transformed with recombinant plasmid encoding HIS-TAN1. After cell pellets were sonicated and centrifuged, the precipitate was dissolved using urea, followed by affinity purification. *E. coli* strain BL21 Star (DE3) was transformed with recombinant plasmid encoding HIS-TAN1-GFP. After cell pellets were sonicated and centrifuged, the precipitate was dissolved using urea, followed by affinity purification (Genscript Corp Piscataway, New Jersey USA). Proteins were refolded and sterilized by filtering. HIS-TAN1 and HIS-TAN1-GFP concentrations were checked with a BCA protein assay (Genscript Corp Piscataway, New Jersey USA). After refolding, HIS-TAN1-GFP was no longer fluorescent. HIS-TAN1-GFP therefore was tagged with an Atto488 dye. HIS-TAN1-GFP was conjugated with Atto488-maleimide (Sigma 28562). 4μM HIS-TAN1-GFP in 80mM PIPES, 1mM MgCl2, 1mM EGTA buffer was reduced with 12.5μM Tris(2-carboxyethyl)phosphine hydrochloride for 10 minutes followed by a 4 hour incubation with 250μM Atto488 dissolved in DMSO (10mM) at room temperature. Unreacted excess dye was removed by running the sample through a 10DG desalting column (BioRad 732-2010) and concentrating with a 30K MWCO PES concentrator (Thermo 88521). HIS-TAN1-GFP and HIS-TAN1-GFP-Atto488 (∼80% degree of labeling) activity was confirmed by microtubule co-sedimentation assay. Conjugation of Atto488 dye was determined by imaging the results of the microtubule cosedimentation assay on a SDS-PAGE experiment using a UV light source showing fluorescent bands corresponding to a Atto488 tagged HIS-TAN1-GFP.

### Microtubule binding and co-sedimentation

A microtubule binding assay kit was used to assess HIS-TAN1 microtubule binding in relation to positive and negative controls, according to manufacturer conditions (Cytoskeleton Inc., MK029). For determining affinity of HIS-TAN1 to microtubules, microtubules were polymerized from 50μM starting concentration of tubulin in the presence of 1mM GTP for 2 hours at 37°C followed by the addition of 10μM taxol. HIS-TAN1 and microtubules were incubated for 25 minutes and spun down at 39,000 x *g* at 25°C. HIS-TAN1-GFP and HIS-TAN1-GFP-Atto488 protein was incubated with microtubules at room temperature for 25 minutes and spun down at 21,000 x *g* at 25°C. Equal amounts of protein samples were loaded into a SDS PAGE experiment (10% gel), stained with Coomassie and analyzed using ImageJ Gel Analysis tool with correction applied due to some HIS-TAN1-GFP-ATTO488 precipitation during the assay in samples without microtubules. Spindowns were performed at least three times for each concentration tested. Statistical analysis was performed using Graphpad Prism. To assess microtubule binding by microscopy, rhodamine labeled microtubules (1:25 rhodamine tubulin:unlabelled tubulin) were polymerized from 50μM starting concentration of tubulin in the presence of 1mM GTP for 2 hours at 37°C followed by the addition of 10μM taxol. 100nM rhodamine labelled microtubules were incubated with 50nM HIS-TAN1-Atto488 for 5 minutes and then pipetted onto a coverslip and imaged.

### Reconstitution of in vitro microtubule dynamics

In-vitro microtubule dynamics were conducted according to previous protocols (Dixit and Ross, 2010). Flow chambers were assembled using silanized coverslips and double-sided sticky tape with a chamber volume of ∼20μL. A 20% monoclonal anti-tubulin antibody (clone BN-34, Sigma, St. Louis, MO) was used to coat the surface followed by blocking with 5% pluronic F-127 (Sigma #P2443) for five minutes each step. Rhodamine and biotinylated guanosine-5′-(α,β-methylene)triphosphate (GMPCPP) microtubule seeds were then flowed into the cell. Microtubule growth was initiated using 17.5μM 1:25 rhodamine-labeled bovine tubulin in 80mM PIPES, 1mM MgCl2, 1mM EGTA with 0.15% methylcellulose (w/v), 100mM DTT, oxygen scavengers (250μg/mL glucose oxidase, 25μg/mL catalase), 5mg/mL glucose, 2mM GTP along with the specified amount of HIS-TAN1 protein. To assess microtubule bundling, a higher concentration of tubulin (22.5μM, 1:25 rhodamine tubulin:unlabelled tubulin) was used in the reaction to promote microtubule growth and crossovers. At least two slides were prepared for each concentration and experimental condition. The samples were excited with a 561-nm (at 4 mW output) diode-pumped solid-state laser (Melles Griot, Albuquerque, NM) and visualized through a 100X objective (NA 1.45) and back-illuminated electron-multiplying CCD camera with a 582-636nm emission filter set using TIRF (ImageEM, Hammamatsu). Images were collected every 2 seconds. Kymographs were used to analyze data in FIJI (Schindelin et al., 2012).

### Predicting Division Planes from Wild-Type and *tan1* Cell Shapes using Surface Evolver

Samples from WT and *tan1* mutant maize plants expressing YFP-TUBULIN (α-tubulin fused to the Citrine variant of Yellow Fluorescent Protein, (Mohanty et al., 2009)) were dissected to the symmetrically dividing leaf zones to identify PPB location. To identify the cell outlines for three-dimensional reconstruction, samples were either stained with 0.1mM propidium iodide or expressed PLASMA MEMBRANE INTRINSIC PROTEIN2-1 fused to CFP to outline the plasma membranes (Mohanty et al., 2009). Three-dimensional cell shape reconstructions were generated using MorphoGraphX, while three-dimensional PPB reconstructions were generated using Trainable Weka Segmentation (Barbier de Reuille et al., 2015; Arganda-Carreras et al., 2017). Cells were collected from more than three individual plants for each genotype. A previous protocol was followed for modeling symmetric divisions by soap-film minimization using Surface Evolver (Brakke, 1992; Martinez et al., 2018). Briefly, cell outlines were smoothed using 30th degree spherical harmonics followed by surface area minimization from 241 starting planes with normals uniformly distributed over a sphere. For PPB offset measurements, the distance between the midplane of the PPB and the surface of the predicted division was measured in microns squared. Abnormality index was defined by the distance of the area surface center and the volume center for the cell. The Surface Evolver based pipeline used is hosted on Github (https://github.com/jdhayes/predictive_division/).

### Colocalization analysis

Maize plants were dissected to reveal the symmetrically dividing leaf zones to image TAN1-YFP and CFP-TUBULIN at the cortex of maize epidermal cells during telophase using a Zeiss 880 LSM. Airyscan super resolution mode was used and the images were processed using default settings. Three separate plants were imaged for the collection of cells. Micrographs were imported into FIJI and cropped to the cell of interest where colocalization was assessed. Just Another Colocalization Plugin (JACoP) was used in order to determine the Pearson Correlation Coefficient and Manders Overlap Coefficient for each cell (Bolte and Cordelières, 2006). Data generated was analyzed using GraphPad (Prism).

### Microscopy for in vitro and in vivo imaging

Taxol stabilized rhodamine labeled microtubules and HIS-TAN1-GFP-Atto488 were visualized on an inverted Nikon Ti stand (Nikon) with a W1 spinning disk (Yokogawa) and a motorized stage (ASI Piezo) run with Micromanager software (micromanager.org) and built by Solamere Technology. Solid-state lasers (Obis) and emission filters (Chroma Technology) used had excitation 561 nm; emission, 620/60 nm (for rhodamine-tubulin); and excitation, 488 nm; emission, 520/50 nm(for HIS-TAN1-GFP-Atto488). A 100x oil lens (1.45 numerical aperture) and Immersion Oil Type FF (Cargille immersion oil, 16212) was used. Maize epidermal cells used for modeling were visualized using a 60× water-immersion objectives with 1.2 numerical aperture. An excitation of 561; emission, 620/60 (for propidium iodide) and excitation of 514; emission, 540/30 (for YFP-TUBULIN). Perfluorocarbon immersion liquid (RIAAA-678; Cargille) was used on the objective.

Dynamic rhodamine-labeled microtubules were excited with a 561-nm (at 4 mW output) diode-pumped solid-state laser (Melles Griot, Albuquerque, NM) using a 100X (NA 1.45) objective and TIRF microscopy. Images were acquired with a back-illuminated electron-multiplying CCD camera (Hamamatsu, Bridgewater, NJ, ImageEM) and rhodamine filter sets (582–636 nm emission).

Colocalization data on TAN1-YFP and CFP-TUBULIN in Figure 4 was collected using a Zeiss LSM 880 Elyra, Axio Observer and a 100x/1.46 Oil lens (Cargille immersion oil, 16212). TAN1-YFP was excited with 514 while CFP-TUBULIN was excited using 458 and imaged using super resolution airyscan mode with a MBS 458/514 and 420-480 BP + LP 605 filter set. Airyscan images were processed using default settings using Zen Black software (Zeiss). FIJI was used for colocalization analysis.

## Supplemental Material

Supplemental Figure 1 shows HIS-TAN1-GFP and HIS-TAN1-GFP-Atto488 microtubule binding and affinity using quantitative microtubule cosedimentation assay.

Supplemental Movie 1 shows HIS-TAN1 mediated microtubule crosslinking events observed during in vitro dynamic microtubule reconstitution assays imaged using TIRF microscopy.

Supplemental Movie 2 shows potential microtubule interactions between the phragmoplast leading edge and TAN1-YFP protein localized at the cortical division site in maize epidermal leaf cells.

## Acknowledgements

Thanks to Ms. Jocelyne Aranda, Mr. Christopher Hoyt, and Ms. Sukhmani Sidhu for collecting some images used for the modeling work presented. We thank Dr. David Carter (University of California, Riverside) for Zeiss LSM 880 training. We also thank the University of California, Riverside Agricultural Operations for greenhouse and field space. Funding from NSF-MCB1716972 and USDA-NIFA to CGR and from NSF-MCB1453726 to RD is gratefully acknowledged. Funding from a Ford Foundation Dissertation Fellowship to PM is gratefully acknowledged.

## Author contributions

CGR, RD, SEO’L and KAB provided equipment, reagents and experimental guidance. PM and RB performed in-vitro experiments with RB offering assistance on microtubule co-sedimentation. PM captured images used for modeling and performed live cell time-lapse. CGR and RD supervised experiments. PM analyzed data and made figures, PM and CGR wrote manuscript with comments and edits from coauthors. CGR and RD acquired funding.

**Supplementary Figure 1.**
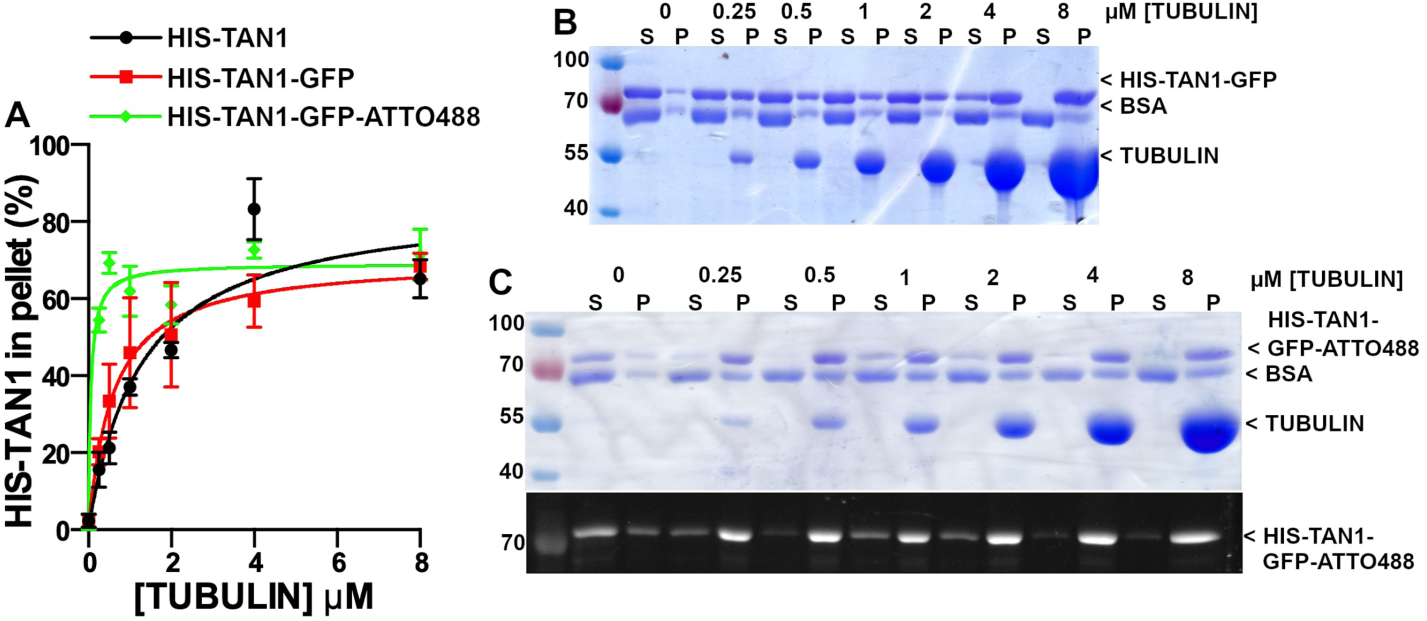
HIS-TAN1-GFP and HIS-TAN1-GFP-Atto488 binding affinity to taxol stabilized microtubules. (A) Co-sedimentation binding data with fits to hyperbolic binding isotherms for HIS-TAN1 (replotted from Figure 1C), HIS-TAN1-GFP, and HIS-TAN1-GFP-Atto488. Apparent affinity for HIS-TAN1-GFP is 0.59μM +/- 0.25, while HIS-TAN1-GFP-Atto488 is 0.06μM +/- 0.03 corrected for the average pelleting in samples without microtubules added (average +/- S.D.) (B) Coomassie stained SDS PAGE experiment from spindown of HIS-TAN1-GFP in the presence of varying concentrations of tubulin (0-8μM). (C) Coomassie stained SDS PAGE experiment from spindown of HIS-TAN1-GFP-Atto488 in the presence of varying concentrations of tubulin (0-8μM). Below Coomassie stained SDS-PAGE experiment, HIS-TAN1-GFP-Atto488 was excited using ultraviolet light source to confirm Atto488 maleimide conjugation with HIS-TAN1-GFP used in the spin-down assays.

Supplemental Movie 1. In vitro microtubule interactions mediated by HIS-TAN1. 2μM HIS-TAN1 was added to a dynamic microtubule assay imaged using TIRF. Arrowheads indicate sites of TAN1 mediated microtubule pulling behavior.

https://drive.google.com/open?id=1jUe-EuH1vxPMJwn7bPWB04Uc09GxcqyJ

Supplemental Movie 2. Phragmoplast leading edge microtubules (CFP-TUBULIN, green) interactions with TAN1-YFP (magenta) at the division site. Arrowheads indicate sites of potential interactions.

https://drive.google.com/open?id=1RdfGC1KpU0nKHp8SKQv4Cq1UGmwGJAUI

